# Acute transient cognitive dysfunction and acute brain injury induced by systemic inflammation occur by dissociable IL-1-dependent mechanisms

**DOI:** 10.1101/127084

**Authors:** Donal T. Skelly, Éadaoin W. Griffin, Carol L. Murray, Sarah Harney, Conor O’Boyle, Edel Hennessy, J Nicholas Rawlins, David M. Bannerman, Colm Cunningham

## Abstract

Systemic inflammation can impair cognition with relevance to dementia, delirium and post-operative cognitive dysfunction. Acute episodes of delirium also contribute significantly to rates of long-term cognitive decline, implying that *de novo* pathology occurs during these acute episodes. Whether systemic inflammation-induced acute dysfunction and acute brain injury occur by overlapping or discrete mechanisms has not been investigated. Here we show that systemic inflammation, induced by bacterial LPS, produces both working memory deficits and acute brain injury in the degenerating brain and that these occur by dissociable IL-1-dependent processes. In normal C57BL/6 mice, LPS (100μg/kg) did not affect working memory but robustly impaired contextual fear conditioning (CFC). However prior hippocampal synaptic loss left mice selectively vulnerable to LPS-induced working memory deficits. Systemically administered IL-1 receptor antagonist (IL-1RA) was protective against, and systemic IL-1β replicated, these working memory deficits. Although LPS-induced deficits still occured in IL-1RI^-/-^ mice, systemic TNF-α was sufficient to induce similar deficits, indicating redundancy among these cytokines. Dexamethasone abolished systemic cytokine synthesis and was protective against working memory deficits despite failing to block brain IL-1β synthesis. Direct application of IL-1β to ex vivo hippocampal slices induced non-synaptic depolarisation and irrevesible loss of membrane potential in CA1 neurons from diseased animals and systemic LPS increased apoptosis in the degenerating brain, in an IL-1RI^-/-^ dependent-fashion. The data suggest that LPS induces working memory dysfunction via circulating IL-1β but dysfunction leading to neuronal death is mediated by hippocampal IL-1β. The data suggest that acute systemic inflammation produces both reversible cognitive deficits, resembling delirium, and acute brain injury that may lead to long-term cognitive impairment but that these events are mechanistically dissociable. This would have significant implications for management of cognitive dysfunction and decline during acute illness.

## Introduction

Peripheral infections are known to trigger episodes of acute cognitive impairment, including delirium, in older populations and in those with dementia (George *et al.*, 1997; Elie *et al.*, 1998). Sterile inflammation, resulting from tissue trauma or surgery, can also induce post-operative cognitive dysfunction and delirium (Kat *et al.*, 2008; van Munster *et al.*, 2008). Cytokines are key mediators of septic and aseptic inflammation and, given its important role in coordinating CNS responses to systemic inflammation (Dantzer *et al.*, 2008; Gosselin & Rivest, 2008), the pro-inflammatory cytokine IL-1β might be predicted equally to underlie infection- and sterile inflammation-induced cognitive dysfunction. Consistent with this idea, IL-1β levels have been associated with delirium in hip fracture patients and in septic encephalopathy (Serantes *et al.*, 2006; Cape *et al.*, 2014b).

Delirium is a profound and acute onset brain dysfunction with impairments in attention and other aspects of cognition, typically triggered by acute medical conditions. Its high prevalence after surgery and infection emphasises the deleterious consequences that systemic inflammation has for cognitive function, particularly in older, cognitively impaired, populations (Davis *et al.*, 2015; Fong *et al.*, 2015). It is now clear that acute systemic inflammation and delirium also significantly increase the risk of long-term cognitive decline and dementia (MacLullich *et al.*, 2009; Davis *et al.*, 2012; Pandharipande *et al.*, 2013) and accelerate the course of existing dementia (Fong *et al.*, 2009; Holmes *et al.*, 2009). Despite the enormous medical and economic implications (Leslie *et al.*, 2008), the question of whether systemic inflammation-induced acute dysfunction and acute brain injury, leading to more persistent deficits, occur by overlapping or discrete mechanisms has not been investigated.

It is well established that IL-1 impacts on cognitive function (Cunningham & Sanderson, 2008; Yirmiya & Goshen, 2011). IL-1 is reported to disrupt consolidation of context-associated fear memory (Barrientos *et al.*, 2002; Goshen *et al.*, 2007; Cibelli *et al.*, 2010) and the IL-1 receptor antagonist, IL-1RA, is protective against post-acquisition contextual fear conditioning (CFC) deficits induced both by systemic LPS- and post-operatively-induced inflammation (Cibelli *et al.*, 2010; Terrando *et al.*, 2010; Barrientos *et al.*, 2012). However, it is likely that systemic inflammation impacts upon multiple cognitive processes. Furthermore, long-term memory consolidation is manifestly different from the acute fluctuating short-term memory processes that are affected in delirium and in relevant models thereof (Brown *et al.*, 2011; Davis *et al.*, 2015) and different from acute neuronal death events described after systemic inflammation (Cunningham *et al.*, 2005). Given the massive public health burden of dementia and delirium, both of which have been associated with IL-1 (Holmes *et al.*, 2003; Heneka *et al.*, 2013; Cape *et al.*, 2014a), it is important to characterise the roles of IL-1β in these different processes. In the current study we hypothesised that 1) bacterial endotoxin (LPS; lipopolysaccharide) would have multiple effects on cognition and neuronal intergrity and that 2) IL-1 would be causative in these changes.

## Materials and Methods

### Animals

Female C57BL/6 mice at 8-12 weeks of age (Harlan Olac Ltd, UK) were housed in cages of 5 at 21°C with a 12:12 h light-dark cycle with food and water *ad libitum.* To test the effects of LPS in cognitive function in young healthy mice (2-4 months) C57BL/6 mice were injected intraperitoneally (i.p.) with 100 (or 200) μg/kg of LPS (*Salmonella Equine abortus*, Sigma (L5886)) in sterile saline.

### Animals with neurodegenerative pathology (stereotaxic surgery)

To examine effects of LPS on cognitive function in animals with prior synaptic loss mice were first intrahippocampally inoculated with 1μl of 10% w/v prion infected-(ME7 strain) or normal brain homogenate (NBH), at co-ordinates from bregma: anterior-posterior -2.0 mm, lateral -1.7 mm, depth -1.6 mm) using a Hamilton microsyringe, under anaesthesia with intra-peritoneal 2,2,2-tribromoethanol. Experimental groups with ME7 or NBH were then injected intraperitoneally (i.p.) with 100 μg/kg of LPS (*Salmonella Equine abortus*, Sigma (L5886)) at 15-16 weeks post-inoculation. LPS produces acute working-memory deficits in ME7 animals at this disease-stage (Murray *et al.*, 2012). Control animals were administered non-pyrogenic saline. In further experiments alternative inflammatory stimuli were examined: IL-1β (R&D Systems, Minneapolis, MN, USA) was injected, i.p. at 15 μg/kg and TNF-α (Peprotech, Rocky Hill, NJ, USA) at 50 μg/kg i.p. in sterile saline.

In experiments using anti-inflammatory interventions IL-1RA (10 mg/kg, i.p.; Kineret, Biovitrum, Sweden) was given immediately before LPS or IL-1β (15 μg/kg); dexamethasone-21-phosphate was administered i.p. at a dose of 2 mg/kg, 60 minutes before LPS. This dexamethasone-21-phosphate dose was sufficient to robustly suppress the systemic secretion of IL-1β, TNF-α and IL-6 (Teeling *et al.*, 2010).

IL-1RI^-/-^ mice (B6.129S7-Il1r1<tm1Imx>; kindly provided by Prof. Kingston Mills) have a null mutation in *Il1r1* and were backcrossed 7 times to C57BL/6 before subsequent maintenance as an inbred colony. C57BL/6 mice were used as control in these experiments. We have detected no significant differences between IL-1RI^-/-^ and C57BL/6 mice in working memory or contextual fear conditioning tasks (Murray *et al.*, 2013). Body temperature was measured using a rectal probe (TH-5 thermoprobe, Physitemp, NJ) 18 hours post-challenge with LPS (750 μg/kg i.p.) in WT and IL-1RI^-/-^ mice inoculated with ME7 or NBH.

All animal procedures were performed in accordance with Irish Department of Health & Children and UK Home Office regulations and all efforts were made to minimize suffering to the animals.

### Working memory: Food-rewarded and escape from water T-maze alternation tasks

We assessed short term/working memory (3-9 hours post-LPS and 1-7 hours post-IL- 1β) using alternation behaviour in both food-rewarded and escape-from-water-motivated T-maze tasks. We had specifically designed an ‘escape from shallow water’ T-maze task to allow assessment of working memory performance in animals experiencing sickness behaviour, as previously described (Murray *et al.*, 2012). The T-maze was constructed of black perspex with the following dimensions (cm): long axis 67, short axis 38, depth 20 and arm width 7. There was a single 40 mm diameter hole at the end of each choice arm, 2 cm from the floor. Black exit tubes were inserted into these holes (these may also be blocked to prevent exit). A ‘guillotine’ door was inserted at the entrance to the choice arms to prevent access to one or other choice arm. This maze was filled with water at 20°C to a depth of 2 cm to motivate mice to leave the maze by “paddling” or walking “on tip-toe” to an exit tube.

Animals were taken with their cage mates to a holding cage. Each mouse was placed, individually, in the start arm of the maze with one arm blocked such that they were forced to make a left (or right) turn, predetermined by a pseudo-random sequence (equal numbers of left and right turns, no more than 2 consecutive runs to the same arm). On making this turn the mouse could escape from the water by entering the small tube, and then a transit tube, in which it was carried to another holding cage. The mouse was held here for 25 seconds (intra-trial interval) during which time the guillotine door was removed and the exit tube was switched to the alternate arm. The mouse was then replaced in the start arm and could choose either arm. The mouse must alternate (i.e. non-match to place) from its original exit arm to escape. On choosing correctly mice escape to the transit tube as before and are returned to their home cage. On choosing incorrectly the mice were allowed to self-correct to find the correct exit arm. Animals were trained for sessions of ten trials (inter-trial interval of 20 minutes) until reaching the criterion of performance of 70% or better (≥ 2 consecutive days), and not showing any evidence of a side preference (i.e. all errors occuring on the same side might indicate a preference for a particular body turn). This level of performance was maintained by running animals 3 times per week until reaching 15 to 16 weeks post-inoculation. At this time, those animals still at criterion (on average about 5% of animals fail to reach criterion) were challenged with LPS, IL-1β (15 μg/kg), TNF-α (50 μg/kg) or saline ± anti-inflammatory interventions.

Naive animals were also trained on an appetitively-motivated version of this task using an enclosed T-maze of the same dimensions. Mice were rewarded on sample and choice runs with 70μl of sweetened condensed milk contained in the lid of a 1.5 ml Eppendorf tube, placed at the end of the rewarded arm. Animals were food-deprived to approximately 90-95 *%* of free-feeding weight for this experiment. Animals choosing the correct arm were allowed to consume the entire reward while those choosing the incorrect arm were removed from the maze without reward and returned to the home cage. After successful baseline performance at above 80% alternation for more than 2 consecutive days, animals were treated with LPS (100 or 200 μg/kg) or with sterile saline and tested for 10 trials in a pseudorandomised sequence with equal numbers of left and right turns, as described for the water T-maze above.

### Contextual Fear Conditioning

Contextual fear conditioning (CFC) was recorded using a clear perspex box (40cm x 10cm x 16cm) with a floor containing metal rods connected to a shock generator (UGO Basile, Italy). The mice were placed into the box and allowed to explore for 2 minutes. A tone at 2.9 kHz for 20 seconds was presented, followed by a shock of 0.4 mA for two seconds. After 2 minutes the tone was repeated for 20 seconds and a second shock of 0.4 mA for two seconds was administered. After a further 30 seconds of exploration mice were removed to a holding cage. After 30 seconds in the holding cage saline or LPS (± saline or IL-1RA) were administered before returning the animals to the home cage. Vehicles and treatments were administered in quick succession at discrete intraperitoneal injection sites. Fear conditioning was assessed in the same location 48 hours later for duration of 5 minutes. Freezing was regarded as the complete absence of movement, except those related to respiration, as originally described (Fanselow, 2000). Auditory fear conditioning was also assessed 48h post-fear conditioning. Animals were placed in a different context (a novel empty cage) for 6 minutes and were allowed to explore for 3 minutes (baseline) before presentation of the tone for 20 seconds. The time spent freezing during the final 3 minutes was recorded (ie during and post-tone). Mice were then placed back into home cage after testing was completed.

### ELISA for Cytokines

Under terminal anaesthesia, the thoracic cavity was opened and blood collected in heparinised tubes directly from the right atrium of the heart. Whole blood was centrifuged at 3000 rpm for 15 minutes at 4°C to remove cells and the remaining plasma aliquoted and stored at -20 °C. These samples were then analysed for IL-1β, TNF-α, IL-6, CXCL1 and CCL2. TNF-α, IL-6, CCL2 and CXCL1 were quantified using R&D systems sandwich-type duo set ELISA kits while IL-1β was analysed in both blood and brain using a Quantikine kit (R&D systems, Minneapolis, MN, USA). A standard protocol was followed as previously described (Murray *et al.*, 2011) except for IL-1β, which was as per manufacturer’s instructions with minor modifications. Hippocampal tissue was homogenised in buffer (150mM NaCl, 25mM Tris-HCl, 1% Triton-x; 10% w/v) containing protease inhibitors (Roche) and centrifuged at 14,000rpm for 10 min. Standards ranged from 7.8 to 500 pg/ml. To ensure that all cytokines were reliably quantifiable using the appropriate standard curves, samples for cytokine assays were serially diluted as follows: TNF-α 1/2 and 1/4, IL-6 1/12, 1/36 and 1/108 and CXCL1 1/9, 1/81 and 1/243. Standards were prepared in the range from 8 to 1000 pg/ml and samples were quantified only if the absorbance fell on the linear portion of the standard curve. Blood and brain samples were also assayed for the presence of human IL-1RA using an R&D Systems quantikine assay, performed according to manufacturers’ instructions using human IL-1RA standards in the range 0-2000 pg/ml. Hippocampal/thalamic tissue punches were homogenised in 150 mM NaCl, 25 mM Tris-HCl and 1% Triton X100 at pH 7.4 before centrifugation at 14,000 rpm for 10 minutes. Supernatants were diluted 1 in 2 in assay diluent in wells pre-coated with anti-human IL-1RA polyclonal antibody.

### RNA isolation, cDNA synthesis and quantitative PCR

The isolation of total RNA, the synthesis of cDNA and analysis of transcription by quantitative PCR have been performed essentially as previously described (Cunningham *et al.*, 2007). Briefly, treated animals were transcardially perfused with heparinised saline and the hippocampus and dorsal thalamus were punched out of coronal brain sections of approximately 2mm thickness. Tissue was snap frozen in liquid nitrogen and stored at -80°C. Total RNA was extracted from brain samples using Qiagen RNeasy Plus^™^ mini kits (Qiagen, Crawley, UK) according to manufacturer’s instructions. Contaminating gDNA was removed using the Qiagen RNase-free DNase I enzyme. RNA yields were determined by spectrophotometry at 260 and 280 nm and stored at –80°C until cDNA synthesis and PCR assay. Using a High Capacity cDNA Reverse Transcriptase Kit (Applied Biosystems, Warrington, UK), cDNA was synthesized from total RNA. 200 ng of total RNA were reverse transcribed in a 10 μl reaction volume and 1 μl of the reverse transcription reaction (RT) was used for polymerase chain reaction (PCR). Reagents were supplied by Applied Biosystems (SYBR^®^ Green PCR Master Mix) and Roche (FastStart Universal Probe Master [Rox], Lewes, UK). Assays were designed using the published sequences for the genes of interest and Primer Express software and primer pairs were checked for specificity by standard RT-PCR using Promega PCR reagents (Promega, Southampton, UK). Primer and probe sequences for those mRNA species have been previously published. Assays were quantified using a relative standard curve, as previously described (Cunningham *et al.*, 2007) constructed from cDNA that was synthesised from 1 μg total RNA, which had been isolated from mice showing up-regulation of all target transcripts of interest in this study.

### Electrophysiology

Transverse hippocampal slices (300 mm) were prepared from the brains of NBH or ME7 mice at 18-19 weeks post-inoculation and from IL-1RI^-/-^ mice at 8 months of age. Slices were cut in ice-cold artificial cerebrospinal fluid (aCSF) solution containing (in mM) 75 sucrose, 87 NaCl, 25 NaHCO_3_, 2.5 KCl, 1.25 NaH_2_PO_4_, 0.5 CaCl_2_, 7 MgCl_2_, 10 D-glucose, 1 ascorbic acid and 3 pyruvic acid. During incubation and experiments slices were perfused with ACSF containing (in mM) 125 NaCl, 25 NaHCO_3_, 2.5 KCl, 1.25 NaH_2_PO_4_, 2 CaCl_2_, 1 MgSO_4_, 25 D-glucose. The slices were maintained at 33 °C for 1 hr following dissection and all recordings were performed at physiological temperature (32-34°C). Whole-cell patch-clamp recordings were made from CA1 pyramidal neurons, visualised using an upright microscope (Olympus BX51 WI, Middlesex, UK) with infra-red differential interference contrast optics (IR-DIC). Patch pipettes were filled with intracellular solution containing (in mM) 130 KMeSO4, 10 KCl, 0.2 EGTA, 10 HEPES, 20 phosphocreatine, 2 Mg2ATP, 0.3 NaGTP, 5 QX-314, 1 TEA (pH 7.3, 290-300 mOsm). Cells were voltage-clamped at -60 mV and input resistance and membrane capacitance were measured in response to a 10 mV depolarising voltage pulse. During current clamp recordings, current injection was initially set to maintain membrane potential at -60 mV and was not altered for the duration of experiments. Recordings were made using a Multiclamp 700B (Molecular Devices, Foster City, CA). Signals were filtered at 5 kHz using a 4-pole Bessel filter and were digitized at 10 kHz using a Digidata 1440 analogue-digital interface (Molecular Devices). Data were acquired and analysed using PClamp 10 and Clampfit (Molecular Devices). IL-1β was bath-applied to slices at concentrations of 0.1 ng/ml and 1 ng/ml. In some experiments TNF-α was bath-applied at concentrations up to 20 ng/ml.

### TUNEL immunohistochemistry

Immunohistochemistry for apoptotic cells was performed on formalin-perfused, wax embedded tissue from ME7 animals on wild-type and IL-1R1^-/-^ backgrounds, 18 hours post-LPS (750 μg/kg i.p.) using the “Dead End” TUNEL staining method (Promega, Southampton, UK). Sections were pre-treated with proteinase K (10 μg/ml) for 5 min and labelling was then performed according to the manufacturers instructions. Once apoptotic cells were labeled with fluorescein sections were blocked with 10% normal goat serum for 30 minutes before incubation with biotinylated anti-flourescein antibody (5 μg/ml) and development of the reaction using the Avidin-Biotin-Complex protocol using hydrogen peroxide as substrate and diaminobenzidine as chromagen. TUNEL-positive cells also showing evidence of nuclear condensation were counted in 10 μm coronal sections of ME7 animals treated with LPS in wild type and IL-1R1^-/-^ mice. At least two sections were counted, in n=5 animals per group, by an observer blind to the experimental conditions.

### Statistical analyses

Behavioural data were compared by repeated measures ANOVA with Bonferroni post-hoc tests performed after significant main effects. Molecular, electrophysiology and TUNEL data were analysed by one or two-way ANOVA as appropriate, while temperature data were analysed by three-way ANOVA, followed by Bonferroni post-hoc tests for a priori selected pairwise comparisons.

## Results

### Differential effects of LPS on different hippocampal-dependent cognitive tasks

We assessed working memory using T-maze alternation tasks that require both attention to, and retention in the working memory of (for 25 seconds), the prior location of an exit in order to determine the new exit location. LPS did not impair working memory in a food-rewarded T-maze alternation protocol at 3-7 hours post-treatment at either 100 or 200 μg/kg i.p. (Fig. 1a). To validate this finding in an aversively motivated working memory task, not reliant on motivation for appetitive rewards, we also used an ‘escape from shallow water’ T-maze working memory task specifically adapted for use in animals experiencing acute sickness behaviour. LPS (100 μg/kg) had no impact on working memory in C57BL6 mice in this T-maze (Fig. 1b).

**Figure 1.**
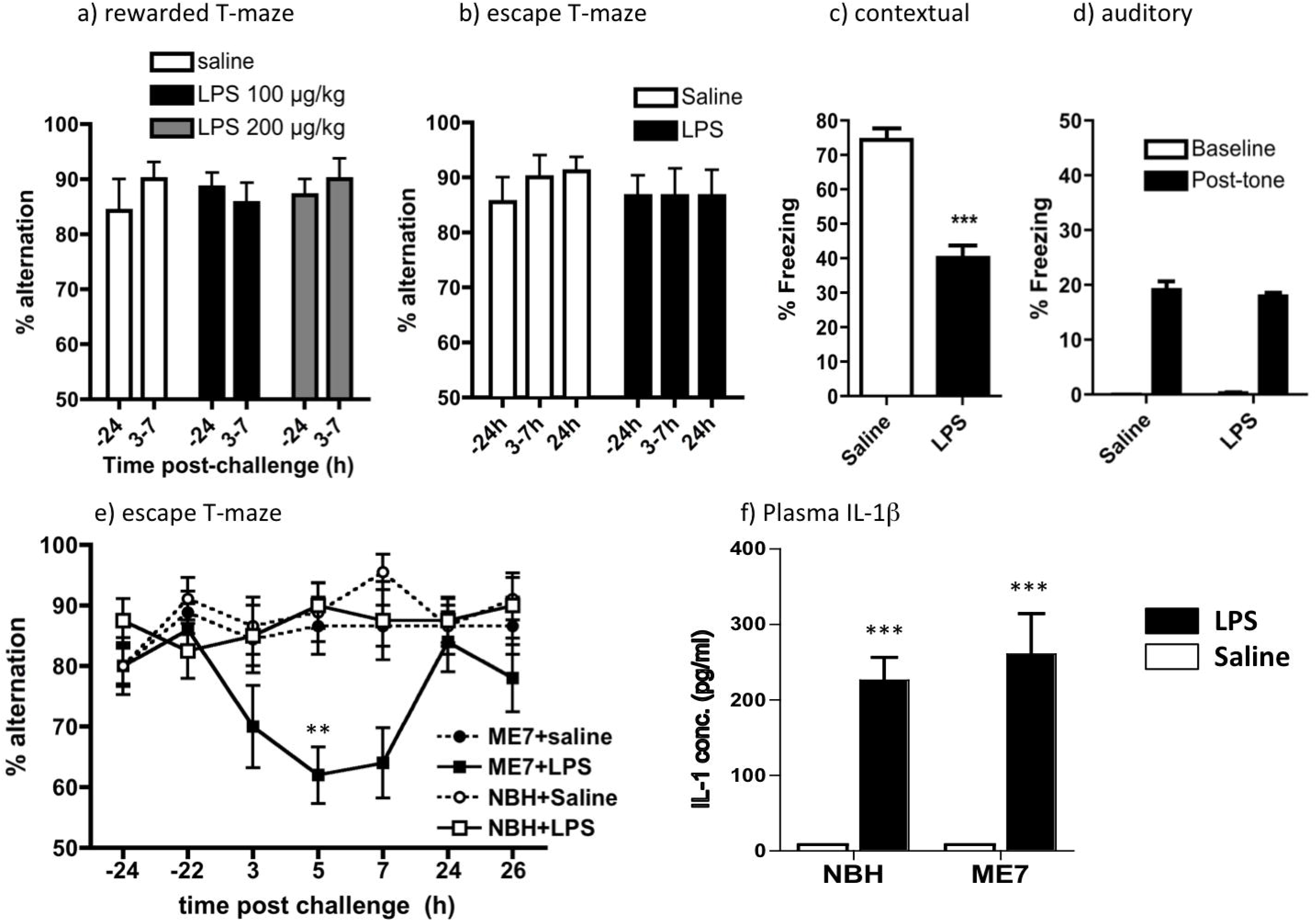
LPS-induced effects on working memory, fear conditioning, and IL-1β expression. Working memory was assessed by **a)** food-rewarded T-maze alternation (all groups n=7) and **b)** escape from shallow water T-maze (both groups n=9) at 24 hours before (-24) and at indicated times post-LPS (100 or 200 μg/kg i.p.). On the X-axis, 3-7 represents % alternation during 10 trials conducted every 20 minutes between 3 and 7 hours post-LPS). Two-way ANOVA analysis found no significant effects of treatment. Contextual **(c)** and auditory **(d)** fear conditioning performance (freezing per 5 minutes) 48 hours post-challenge with LPS (100 μg/kg i.p.). Data are expressed as mean±SEM and analysed by t-test (*** p<0.0001; n=12 for saline and 16 for LPS). **e)** Working memory performance of NBH and ME7 animals 16 weeks post-ME7 inoculation, challenged with LPS (100 μg/kg i.p.) or saline, assessed by T-maze alternation for 10 trials 24 hours pre-challenge, 15 trials between 3-9 hours post-challenge and 10 trials 24 hours post-challenge. Full ANOVA analysis is described in the main text. Significant Bonferroni post-hoc differences between ME7+LPS and both NBH+LPS and ME7+saline are denoted by ** (p<0.01). ME7+LPS n=10, ME7+saline, NBH+saline n=9, NBH+LPS n=8. **f)** Plasma levels of IL-1β, were assessed by ELISA in 16-week ME7 and NBH animals 2 hours post-LPS treatment. Two-way ANOVA revealed a main effect of LPS challenge on circulating IL-1β (F=60.06, df 1,12, p<0.0001) but no main effect of disease and no interaction between treatment and disease (*** p<0.001, denotes the main effect of LPS, all groups n=4). All data are presented as mean ± SEM.

However, LPS (100 μg/kg i.p.) given immediately post-acquisition, was sufficient to significantly decrease freezing in CFC 48 hours after exposure to context and foot-shock pairing (Fig. 1a, p<0.001) but had no impact on auditory fear conditioning (Fig 1d). Therefore, systemic LPS has differential effects on two hippocampal-dependent tasks: disrupting consolidation of contextual memory but not affecting working memory in normal animals.

### LPS differentially affects working memory in vulnerable animals despite equivalent circulating IL-1β □□□□□□

The same LPS challenge was sufficient to produce robust, acute and transient working memory deficits in animals with prior neurodegenerative pathology induced by the ME7 model of prion disease, which peaked at 5 hours and had resolved by 24 hours (Fig 1e), replicating our prior studies (Davis et al., 2015). Repeated measures 3-way ANOVA showed that treatment (LPS) affected performance differently in NBH and ME7 animals (interaction between disease and treatment, F_1,32_=4.64, p=0.0388) and ME7+LPS at it’s nadir (5h) was significantly different to both ME7+saline and NBH+LPS (p<0.01, Bonferroni post-hoc). Therefore spatial working memory remains intact when normal animals are challenged with LPS but is impaired when animals with prior disease are similarly challenged. Despite this selective vulnerability of ME7 animals to peripheral LPS-induced deficits, systemic IL-1β synthesis in ME7 and NBH animals at 2 hours post- LPS (100 μg/kg) was equivalent (Fig. 1f).

### Dissociable effects of IL-1RA treatment on LPS-induced cognitive deficits

Since the acute cognitive impairment observed in 16-week ME7 animals following systemic LPS challenge was immediately preceded by robust circulating IL-1β levels, we assessed whether systemic treatment with the receptor antagonist IL-1RA would be protective against the T-maze and CFC deficits described in figure 1. IL-1RA at 10 mg/kg (i.p.) was sufficient to significantly protect against LPS-induced T-maze deficits in ME7 animals (Fig 2a). There was a significant interaction of treatment and time (F_12,144_=2.4, p=0.0074) and ME7+LPS+IL-1RA and ME7+LPS+veh were significantly different at 5 hours (p<0.01; Bonferroni post-hoc). Both LPS-treated groups were impaired at 7 hours.

**Figure 2.**
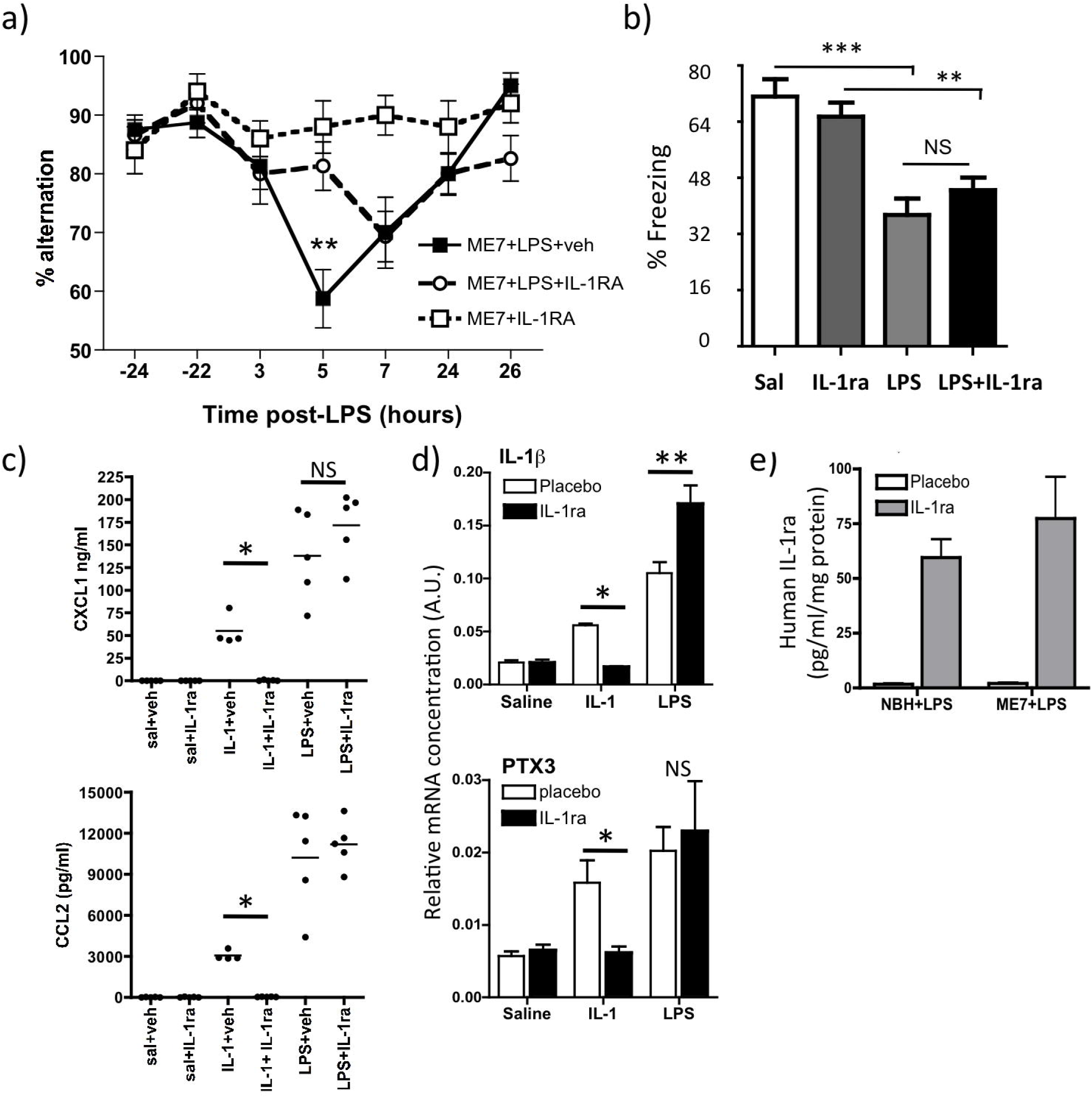
Dissociable effects of systemic IL-1 receptor antagonist (IL-1RA) on T-maze alternation and contextual fear conditioning. **a)** Working memory performance of ME7 animals, 16 weeks post-inoculation, challenged with LPS (100 μg/kg i.p.) in the presence or absence of IL-1RA (10 mg/kg i.p. immediately following LPS). n=17 except ME7+IL-1RA (n=10). Significant Bonferroni post-hoc (p<0.01) after significant 2-way ANOVA denoted by **. **b)** Performance of normal mice in CFC (time spent freezing 48 hours following 0.4 mA foot shock) following systemic challenge with saline or LPS (100μg/kg i.p), in the presence or absence of IL-1RA (10mg/kg i.p). All groups were n=11 except WT+saline (n=7). Data are expressed as mean±SEM and were analysed by two-way ANOVA. Significant differences, assessed by Bonferroni *post-hoc* after significant ANOVA are denoted by ** (p<0.01) and *** (p<0.001). **c)** Effect of i.p. recombinant IL-1RA on plasma chemokine (CXCL1, CCL2) induction 2 hours post-administration of IL-1β (15 μg/kg, i.p.) or LPS (100 μg/kg, i.p.), analysed by ELISA. Bonferroni post-hoc tests (after significant ANOVA) are annotated by * (p<0.05). **d)** Expression of pro-inflammatory genes IL-1β and PTX3, examined in the hippocampus after the same IL-1β or LPS challenges ± IL-1RA. Significant differences by Bonferroni post-hoc test are denoted by *p<0.05 and **p<0.01. All data have been presented as mean±SEM, n=5 per group (except IL-1μ+veh n=4). e) Hippocampal tissue of NBH/ME7 animals treated with LPS in the presence or absence of human IL-1RA, assayed for human IL-1RA by ELISA.

Conversely, systemic IL-1RA at the same dose had no protective effect against the LPS-induced CFC deficit (Fig. 2b). LPS robustly induced decreased freezing in C57BL6 mice (main effect of LPS by two-way ANOVA, F_3,36_=38.68, p<0.0001). Freezing, 48 hours after co-treatment with LPS+IL-1RA (10 mg/kg i.p.), was not significantly different from that induced by LPS (Fig. 2a, F=0.08). Therefore IL-1RA is protective against LPS-induced working memory deficits but not against LPS-induced impairment of consolidation of contextual memory.

Efficacy of i.p. IL-1RA (10 mg/kg) to block systemic and central (hippocampal) inflammatory effects of LPS (100 μg/kg, i.p.) and IL-1β (15 μg/kg, i.p.) was tested. Plasma was prepared 2 hours post-treatment with recombinant IL-1RA (10mg/kg, i.p.) and simultaneous injection of IL-1β, LPS or saline. Both LPS and IL-1β increased plasma CXCL1 and CCL2. IL-1RA had no effect on LPS-induced circulating chemokine but completely blocked IL-1-induced chemokine (p<0.05, one way ANOVA with Bonferroni post-hoc, Fig. 2c).

Hippocampal expression of pro-inflammatory genes IL-1β and PTX3 were also increased by both systemic LPS and IL-1β treatments (Fig. 2d). Once again IL-1- induced increases in both IL-1β and PTX3 were inhibited by systemic IL-1RA (p<0.05), but this treatment failed to block, and infact even appeared to enhance, LPS-induced hippocampal increases in IL-1β and PTX3 (2d). These data indicate that systemic IL-1RA successfully blocks systemic IL-1β actions but LPS still produces plasma and brain inflammatory profiles despite successful inhibition of systemic IL-1β action. Therefore LPS can clearly signal to the brain in the absence of circulating IL-1β actions.

We used an anti-human IL-1RA ELISA to assess whether any of the peripherally applied human IL-1RA reaches the hippocampus, since it could be hypothesised that IL-1RA could show protective effects because of compromised blood brain barrier allowing access of IL-1RA selectively to the ME7 hippocampus. Low levels of human IL-1RA were detectable in the hippocampus of all IL-1RA-injected animals but concentrations were equivalent in NBH and ME7 animals (Fig. 2e). These levels (60-70 pg/ml/mg protein) are almost 7,500-fold lower than plasma levels (481,511 ± 95,460 pg/ml). These data indicate that (i) a tiny fraction of IL-1RA penetrates the brain parenchyma, (ii) that this is not significantly higher in neurodegeneratively diseased (ME7) animals and (iii) that blood levels of IL-1RA remain high at 3 hours post-treatment.

### IL-1β, TNF-α and redundancy in inflammation-induced cognitive dysfunction

To further understand the contribution of IL-1 signalling to the acute working memory deficits induced by LPS in ME7 mice, we inoculated wild type and IL-1R1^-/-^ mice with ME7 and, at 16 weeks into disease progression, challenged these mice with LPS (100 μg/kg i.p.) or saline (Figure 3a). Since LPS produced no working memory deficits in NBH animals (Fig. 1e), NBH controls were not included in these studies on the mechanism of LPS-induced deficits. LPS treatment induced working memory impairments in IL-1R1^-/-^ ME7 animals (treatment effect: F_12,245_=16.61, p<0.0001) and the deficits were not significantly different to those in wild type ME7+LPS animals at any time point (All Bonferroni post-hoc tests p>0.05, Fig. 3a). Thus, IL-1RI is not essential for LPS-induced acute working memory deficits in ME7-inoculated animals. However IL-1RI^-/-^ mice also showed the normal appearance of LPS-induced sickness and weight loss (which was equivalent at 24 hours post-LPS in WT mice and IL-1R1^-/-^ mice: 3.22g vs 2.96g). This is consistent with previous data showing that IL-1R1^-/-^ mice display the full spectrum of innate immune and sickness behavioural responses to LPS (http://jaxmice.jax.org/strain/003245.html) and indicates significant redundancy of specific cytokines in coordination of inflammatory responses to LPS.

**Figure 3.**
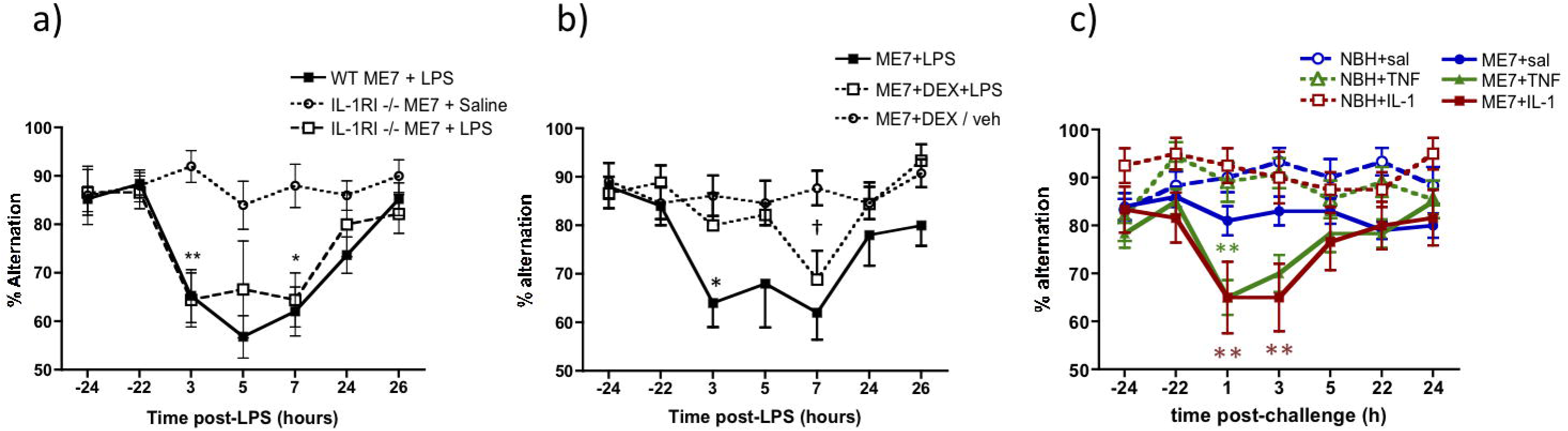
LPS-induced working memory deficits in ME7 animals are intact in IL-1R1^-/-^ mice but are blocked by inhibition of systemic cytokine synthesis and replicated by administration of IL-1β or TNF-α. a) T-maze alternation, of WT and IL-1RI^-/-^ animals, 16 weeks post-inoculation with ME7, in the presence or absence of systemic LPS challenge (100 μg/kg i.p.) was assessed for 10 trials 24 hours before acute challenge, 15 trials 3-9 hours post-challenge and 10 trials 24 hours after the challenge. WT ME7+LPS and IL-1R1^-/-^ ME7+LPS were not different at any time. Significant Bonferroni post-hoc differences between IL-1R1^-/-^ ME7+LPS and IL-1R1^-/-^ ME7+saline are denoted by *p<0.05 and **p<0.01 (n=10 except WT ME7+LPS n=19). b) Working memory performance after systemic LPS (100 μg/kg i.p.) was assessed in the presence or absence of dexamethasone-21-phosphate (2 mg/kg i.p.). Significant post-hoc differences between ME7+LPS+veh and ME7+LPS+DEX (by Bonferroni post-hoc comparisons after a significant ANOVA) are denoted by * (p<0.01) and between ME7+LPS+DEX and ME7+DEX/veh by † (p<0.01). IL-1R1^-/-^ ME7+LPS n=10, IL-1R1^-/-^ ME7+Dex+LPS n=9 and IL-1R1^-/-^ ME7+Dex/vehicle n=13. c) Systemic administration of IL-1β (15 μg/kg) or TNF-α (50 μg/kg) induced acute working memory deficits in T-maze alternation. All animals were assessed for 10 trials 24 hours before acute challenge, 15 trials 1-7 hours post-challenge and 10 trials 24 hours after the challenge. Group sizes were as follows: NBH+saline (n=16), NBH+TNF (n=11), NBH+IL-1β (n=14), ME7+saline (n=20), ME7+TNF (n=12), ME7+IL-1β (n=12). Data pertaining to TNF-α originally published in Hennessy et al., 2017, BBI, 10.1016/j.bbi.2016.09.011(Hennessy *et al.*, 2017). Significant main effects and interactions are described in the main text. Significant differences by Bonferroni post-hoc analysis between ME7+IL-1β and NBH+IL-1β (red) and between ME7+TNF-α and NBH+TNF-α (green) are denoted by ** (p < 0.01). All data are presented as mean±SEM.

Therefore we used dexamethasone-21-phosphate (DEX; 2 mg/kg) to block multiple LPS-induced pro-inflammatory cytokines and to assess its impact on LPS-induced T-maze impairments in IL-1R1^-/-^ ME7 mice (Fig. 3b). This dose was sufficient to block systemic secretion of IL-1β, TNF-α and IL-6 (Fig. S1 and, (Teeling *et al.*, 2010)). DEX alone had no significant impact on T-maze alternation in this paradigm. LPS induced clear working memory impairments in ME7 animals and this was significantly reduced in animals pre-treated with DEX (Fig. 3b). Repeated measures two-way ANOVA showed a main effect of treatment (F_2,30_=8.7, p=0.001) and a treatment x time interaction (F=2.28, df 12,180, p+0.01). ME7+LPS was significantly different to both DEX groups at 3 hrs (Bonferroni post-hoc, p<0.01), although ME7+LPS and ME7+LPS+DEX were no longer significantly different at 7 hours. Thus DEX protects against LPS-induced deficits but some cognitive impairment does eventually occur despite DEX inhibition of systemic cytokine synthesis.

These data suggested that systemic cytokines contribute to LPS-induced working memory impairments. Therefore, we interrogated the roles of specific systemic cytokines. We treated animals with IL-1β (15 μg/kg i.p.) and assessed for working memory deficits. Moreover, it has been demonstrated that in IL-1RI^-/-^ mice, TNF-α compensates for the lack of IL-1 signalling and mediates sickness behavioural and weight loss responses to LPS (Bluthe *et al.*, 2000). Therefore, in a separate set of experiments, the impact of systemically administered TNF-α on T-maze performance was also assessed in both normal (NBH) and ME7 animals. Impairments were not produced by either cytokine in NBH animals but both IL-1β and TNF-α challenges induced acute and transient working memory deficits in ME7 animals (Figure 3c). The IL-1β experiment showed an interaction of disease and treatment (F_1,53_=5.42, p=0.0236) and ME7+saline and ME7+IL-1β animals were significantly different at 1 and 3 hours (p<0.05). As previously described (Hennessy *et al.*, 2017), Bonferroni post-hoc analysis showed that ME7+TNF-α animals were significantly different from both NBH+TNF-α and from ME7+saline at 1 hour after a significant interaction of disease, treatment and time in 3-way ANOVA analysis (F=3.09, df 6,328, p<0.01). Thus, systemic IL-1β and TNF-α □□□□ cause acute and transient working memory deficits in animals with prior neurodegenerative pathology but have no effect on working memory in normal animals.

### Neuronal sensitivity to IL-1β

Since equivalent systemic IL-1 levels produce differential cognitive outcomes in ME7 and NBH animals, we predicted that neurons of the degenerating brain might be more sensitive to equivalent concentrations of IL-1. To address this hypothesis we applied IL-1β directly to *ex vivo* hippocampal slices and performed whole-cell patch clamp recordings from CA1 pyramidal cells from NBH and ME7 animals at 18-19 weeks post-inoculation.

CA1 pyramidal neurons from ME7 animals had a significantly depolarised resting membrane potential (V_m_) compared to those from NBH animals (-61 ± 2 mV in NBH and -51 ± mV in ME7, n = 13 cells from 5 NBH animals and 24 cells from 8 ME7 animals, unpaired t-test, * p<0.0001, Fig. S2).

We then tested the effects of bath application of IL-1β while recording from cells in current clamp mode, using current injection adjusted to maintain membrane potential at -60 mV during baseline recordings (Fig. 4 a, b, c). Neurons from ME7 animals were significantly more sensitive to IL-1β, which depolarised resting membrane potential by 21 ± 4 mV at 0.1 ng/ml in ME7 animals (n = 13 cells from 5 animals) but had little effect in NBH animals (1.5 ± 2 mV; n=5 cells from 3 animals; p=0.013 by Bonferroni post-hoc test after a significant one-way ANOVA, Fig. 4d). IL-1β also depolarised ME7 animals’ resting membrane potential by 44 ± 7 mV at 1 ng/ml (n = 5 cells from 3 animals) but had milder effects in NBH animals (6 ± 3 mV; n=5 from 3 animals; p<0.001, Bonferroni post-hoc test after significant one-way ANOVA, Fig. 4d). ME7 were thus, significantly more sensitive to IL-1β-induced depolarization by both concentrations of IL-1β tested.

**Figure 4.**
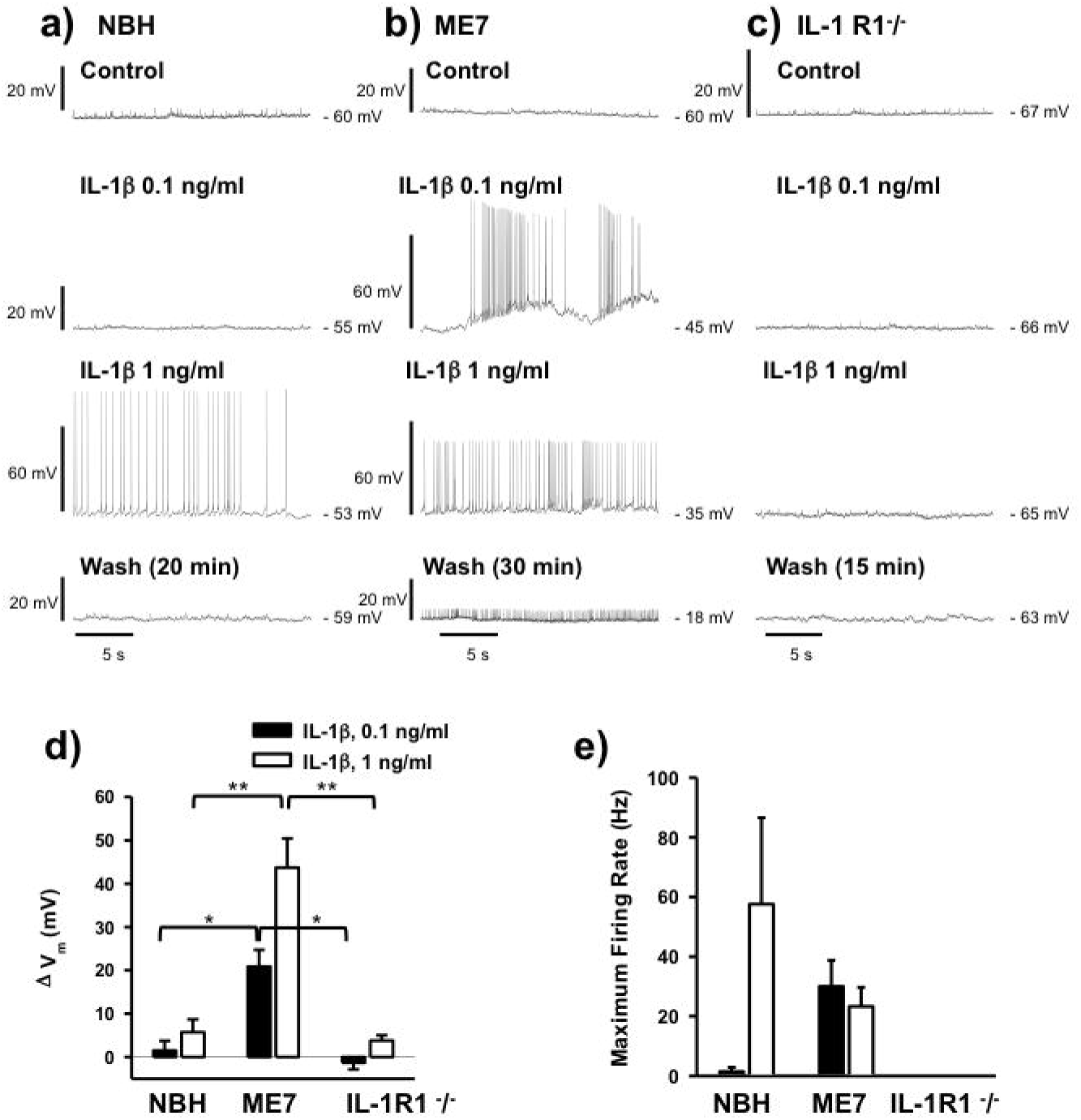
Differential sensitivity of CA1 pyramidal cells to IL-1β in hippocampal slices from NBH and ME7 animals. Traces illustrating current clamp recordings from CA1 pyramidal cells from NBH (a), ME7 (b) and IL-1RI^-/-^ (c) animals during control, baseline recordings and in the presence of IL-1β at concentrations of 0.1 and 1 ng/ml and following washout. The effects of IL-1β were quantified by measuring the change in baseline membrane potential (ΔV_m_), averaged over 10 s, (d) and changes in the maximum action potential firing rate (e). Neurons from ME7 animals were more sensitive to IL-1β, exhibiting greater depolarisation of V_m_ in response to IL-1β at either concentration and a much higher firing rate at the lower concentration of 0.1 ng/ml (n = 5 cells from 3 NBH animals, n = 13 cells from 5 ME7 animals for 0.1 ng/ml and 5 cells from 3 ME7 animals for 1 ng/ml). No action potential firing was induced in cells from IL-1 R1^-^/^-^ animals at either concentration of IL-1β tested (n = 4 cells from 2 animals). Statistically significant differences between different animal groups treated with 0.1 ng/ml and different animal groups treated with 1 ng/ml are assessed are denoted by *p<0.05 or **p<0.01 (Bonferroni post-hoc tests after significant one way ANOVAs). All data are mean ± SEM.

As a consequence, IL-1β induced action potential firing in ME7 CA1 neurons at 0.1 ng/ml, with a maximum firing rate of 30 ± 9 Hz (0.1 ng/ml, n=13 cells) but did not do so in NBH animals (Fig. 4e). At 1 ng/ml, IL-1β induced spiking at maximum rates of 23±6 Hz (1 ng/ml, n=5 cells, Fig. 4e) in ME7 animals and at a highly variable maximum rate of 58 ± 29 Hz in NBH neurons (n=5 cells, Fig. 4e). The lower firing rates in ME7 cells can be attributed to the greater depolarisation induced by IL-1β, acting to increase sodium channel inactivation and therefore limiting spike firing frequency. Depolarisation of ME7 neurons persisted even after washout of IL-1β, with cells becoming more depolarized to a level where action potential firing was inhibited, suggesting that exposure to IL-1β had an irreversible, detrimental effect on these cells.

The effect of IL-1β on ME7 neurons does not appear to be mediated by modulation of excitatory synaptic transmission since 1 ng/ml IL-1β had no effect on evoked excitatory postsynaptic currents (EPSC amplitude 105 ± 32 % of control amplitude, n = 4 cells from 2 ME7 animals, Supplemental Fig. S3). The ability of IL-1β to induce spiking activity in CA1 neurons was dependent on IL-1R1 expression since IL-1β had little effect on resting membrane potential and did not induce action potential firing in slices from IL-1R1^-^/^-^ animals (n=4 cells from 2 animals, Fig. 4 c, d, e). Moreover, TNF-α (20 ng/ml) had no effect on membrane potential in slices from IL-1R1^-^/^-^ animals (-64 ± 2 mV in control and - 64 ± 2 mV in TNFα, n=5 cells from 2 animals, Supplementary data, S4).

### Consequences of exaggerated IL-1 responsiveness: apoptosis and exaggerated sickness response

Since neurons in the diseased brain were more sensitive to the effects of IL-1 and failed to recover their resting membrane potential, we hypothesised that the previously reported neuronal death induced by systemic LPS (Cunningham *et al.*, 2005) would be mediated by IL-1RI. ME7 and NBH animals were challenged with LPS (750 μg/kg i.p.) and euthanised 18 hours later, the time at which LPS-induced neuronal apoptosis has previously been demonstrated (Cunningham *et al.*, 2005). TUNEL labelling was performed to assess for apoptotic cell death in 10 μm coronal sections. LPS induced robust apoptosis in wild-type ME7 animals, with respect to ME7+saline animals and the number of apoptotic cells was significantly reduced in IL-1R1^-/-^ ME7+LPS animals (Fig. 5 a, b). Two-way ANOVA showed an interaction of treatment and genotype (F=6.05, df 1,16; p=0.026) and Bonferroni post-hoc comparison showed that ME7+LPS was significantly more affected than the ME7 IL-1R1^-/-^+LPS group (p<0.05). Therefore, LPS-induced apoptosis in ME7 animals is partially mediated by IL-1RI.

**Figure 5.**
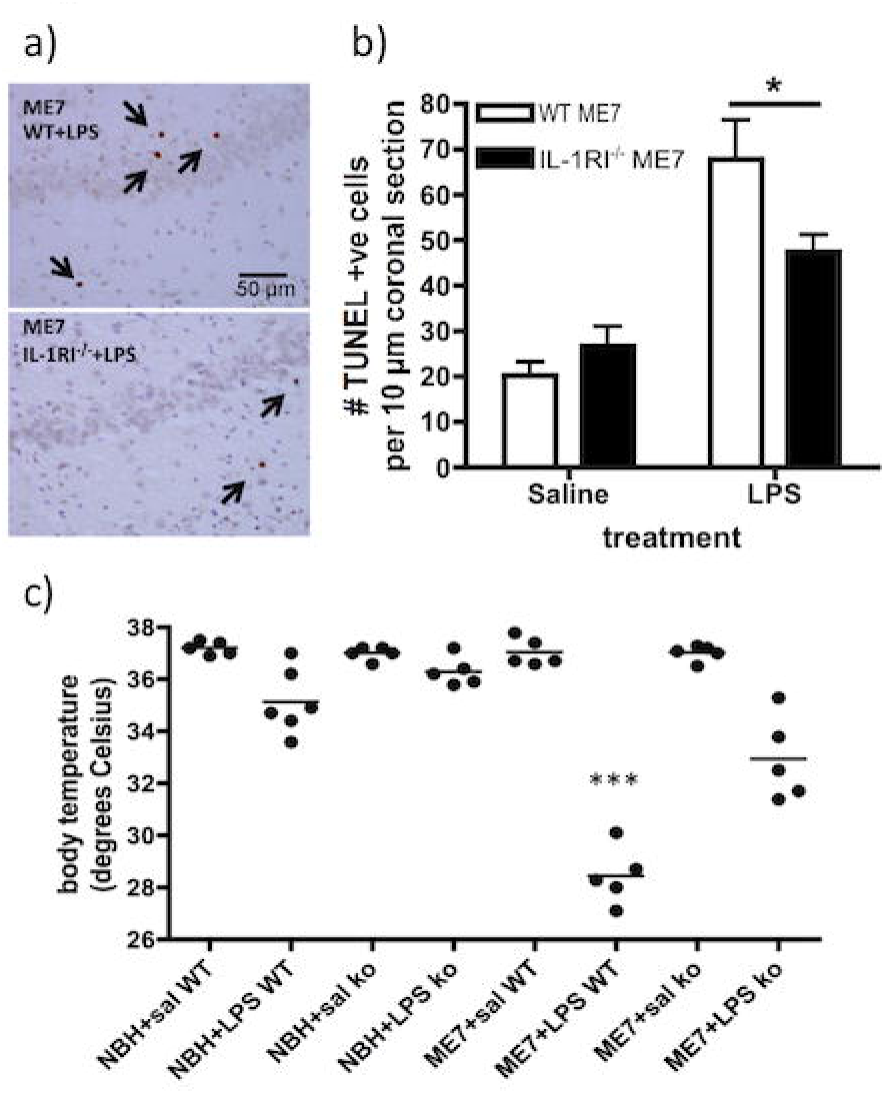
IL-1RI-dependence of LPS-induced apoptosis and exaggerated sickness. ME7 and NBH animals, on a wild-type or IL-1R1^-/-^ background, were challenged i.p. with LPS (750 μg/kg) or saline. a) Representative hippocampal fields after TUNEL immunohistochemistry for apoptotic cells at 18 hours post-LPS or sterile saline, with condensed TUNEL-positive cells labelled with arrows. Scale bar = 50 μm. b) Apoptotic cells were counted in 10 μm coronal sections. Data are shown as mean ± SEM with n=5 in all groups. * denotes ME7+LPS significantly different to ME7 IL-1R1^-/-^+LPS (p<0.005) by Bonferroni post-hoc after significant interaction between strain and treatment by two-way ANOVA (F=6.05, df 1,16; p=0.0257). c) WT and IL-1RI^-/-^ (ko) animals were also assessed (immediately before euthanisation at 18 hours) for core-body temperature. Data are shown as scatter dot plots with the mean displayed. *** denotes ME7+LPS was significantly different both to NBH+LPS and to ME7 IL-1R1^-/-^+LPS (p<0.001) by Bonferroni post-hoc test after significant 3 way ANOVA).

Furthermore, IL-1RI-dependence of the exaggerated CNS response of ME7 animals to systemic LPS was validated by measuring LPS-induced hypothermia. Animals were assessed for core-body temperature directly before euthanisation at 18 hours post-LPS. LPS-treated NBH animals still showed significant hypothermia at this time, but this effect was very much exaggerated in ME7+LPS animals (Fig. 5c; p<0.01). In IL-1R1^-/-^ ME7+LPS animals, this exaggerated effect was very significantly mitigated (p<0.01; all analyses by selected Bonferroni pairwise comparisons after a significant interaction of disease, strain and treatment by 3 way ANOVA: F_1,33_=7.97, p=0.008). Therefore, the exaggerated sickness induced by LPS in ME7 animals is to a substantial extent mediated via IL-1R1.

## Discussion

We have demonstrated that systemic inflammation impacts upon cognitive and neuronal function in multiple ways by dissociable IL-1-dependent mechanisms. While LPS has no impact on working memory in normal animals (at 100 or 200 μg/kg) it robustly impairs working memory in animals with existing neurodegenerative pathology. A systemic-cytokine-mediated mechanism of LPS-induced working memory deficits is supported by systemic IL-1RA and dexamethasone experiments and both systemic IL-1β and TNF-α were sufficient to mimic LPS-induced working memory impairments. Systemic LPS also induced IL-1R1-dependent apoptotic cell death in the neurodegenerating brain and this was consistent with the observation that CA1 neurons in hippocampal slices showed heightened vulnerability to IL-1β-induced loss of membrane potential, which did not recover upon washout. The data suggest that systemic inflammation induces both transient cognitive dysfunction and lasting brain injury and that IL-1 contributes to both processes, but by dissociable mechanisms.

It is now clear that systemic inflammation can trigger episodes of delirium in susceptible individuals (Davis *et al.*, 2015) and that episodes of delirium interact synergistically with existing cognitive impairment to accelerate dementia progression in a manner that is apparently not mediated by exacerbation of classical CERAD markers of dementia (Davis *et al.*, 2017). The implication is that delirium modifies dementia but not via classical pathological hallmarks. Therefore understanding the mechanistic basis of both the acute deficits and the lasting injury is of crucial importance. The current data adds to our knowledge on transient and lasting effects of systemic inflammation on the already degenerating brain.

### Dissociations in inflammation-induced, hippocampal-dependent cognitive impairment

Although impairment of memory consolidation in the CFC paradigm is the best characterised effect of systemic inflammation on cognitive function in young healthy rodents (whether induced by LPS, infection or surgical trauma) (Goshen *et al.*, 2007; Cunningham & Sanderson, 2008; Cibelli *et al.*, 2010), we have argued that this consolidation of long-term memory may not be optimal for studying the dynamic and fluctuating attentional and working memory deficits observed during delirium (Cunningham & Maclullich, 2013). Hippocampal IL-1β clearly impairs late-phase LTP (Tong *et al.*, 2012) and impairs consolidation of contextual memory both in young and aged LPS*/E*. coli-treated rats (Frank *et al.*, 2010) and in post-operative mice (Cibelli *et al.*, 2010). However, the consolidation of memory that is central to CFC is associated with impairment in late-phase LTP (Chapman *et al.*, 2010) and deficits in this relatively slow, protein synthesis-dependent, process cannot explain the disruption in dynamic short-term memory and/or attentional processes described here. There is no intuitive role for memory consolidation in the current T-maze task and the neurological substrates required for CFC and T-maze alternation are manifestly different (Bannerman *et al.*, 2014). Here we show that working memory and contextual memory consolidation are differently affected by systemic inflammation: despite the CFC impairment observed with 100 μg/kg LPS in the current study, neither this nor 200 μg/kg LPS was sufficient to impair working memory in the T-maze in normal animals. Therefore systemic inflammation differentially affects different hippocampal-dependent tasks, as has previously been suggested from context discrimination tasks, (Czerniawski & Guzowski, 2014; Czerniawski *et al.*, 2015).

However the same i.p. LPS dose that failed to impair working memory in normal animals was sufficient to produce robust working memory deficits in animals with prior hippocampal synaptic loss (see (Davis *et al.*, 2015)). These deficits were ameliorated by peripheral treatment with IL-1RA, but this treatment had no beneficial effect with respect to the CFC deficits already described. These data illustrate that systemic inflammation produces impairments in multiple cognitive domains and that, at least with respect to these 2 hippocampal-dependent tasks, there are dissociable IL-1-dependent mechanisms. IL-1β is described to act centrally to impair consolidation in LPS-induced CFC deficits (Barrientos *et al.*, 2002; Goshen *et al.*, 2007; Cibelli *et al.*, 2010; Terrando *et al.*, 2010; Barrientos *et al.*, 2012), but appears to act systemically here to impair working memory, as we discuss below.

### Systemic versus central effects

That the working memory impairment induced by LPS is mediated by circulating rather than centrally-produced IL-1β is supported by multiple strands of evidence. 1) Peripherally applied IL-1RA was protective against the working memory deficits and CNS concentrations were 1/7500 of levels in the plasma at the time of protection against the deficits, consistent with prior data that peripherally administered IL-1RA (17 kDa) can only cross the blood brain barrier (BBB) to a limited extent (Gutierrez *et al.*, 1994). Moreover, although IL-1RA entry to the brain is thought to occur primarily when and where the BBB has been breached (Greenhalgh *et al.*, 2010) we observed similar levels in NBH and ME7 animals arguing against the idea that existing neurodegenerative disease might have made the brain more permeable to IL-1RA 2) Cytokine and chemokine analysis shows that IL-1RA blocked IL-1 action in the blood (Fig. 2c) but did not prevent LPS-induced changes in hippocampal inflammatory transcripts, including *de novo* IL-1. This is consistent with data showing direct activation of the brain vasculature by systemic LPS (Singh & Jiang, 2004) and CNS inflammatory mediator production occurring despite abrogation of systemic cytokines via dexamethasone-21-phosphate inhibition (Murray *et al.*, 2011) or by depletion of peripheral TLR4-positive macrophages (Chakravarty & Herkenham, 2005; Gosselin & Rivest, 2008; Serrats *et al.*, 2010; Chen *et al.*, 2012). 3) The DEX regime used here was 90% effective in reducing systemic IL-1β synthesis but not at all effective at blocking CNS transcription and microglial synthesis of IL-1β (Fig. S1 and (Murray *et al.*, 2011)). 4) Systemic administration of IL-1β is sufficient to induce the same deficits as LPS and the impairments in performance are apparent by 1 hour post-injection, too soon to have been the result of induced *de novo* brain IL-1β synthesis. Thus IL-1RA appears to act peripherally rather than centrally to block IL-1 action with respect to working memory.

### Time-dependent protection and redundancy in inflammation-induced impairments

The protection afforded by IL-1RA is temporary (Fig. 2a). IL-1RA remained at 481.5 ng/ml at 3 hours post-injection in the current study, compared to 125 pg/ml plasma IL-1β 2h post LPS at 100 μg/kg (Skelly *et al.*, 2013), but rapid renal metabolism of IL-1RA (Cawthorne *et al.*, 2011), may cause levels to fall below therapeutic efficacy (Barrientos *et al.*, 2012) during the later trials (7-9 h post-LPS). The protection offered by dexamethasone administration in IL-1R1^-/-^ animals also waned at 7h post-challenge. Given that CNS inflammatory mediator production persisted even in the presence of systemic IL-1RA or dexamethasone (Fig. S1), it is plausible that propagation of central mediators such as IL-1β may have additional effects, or expression of additional inflammatory mediators sufficient to disrupt cognition, may also have occurred independent of systemic IL-1β. One such candidate, which may contribute to systemic IL-1-independent effects, is TNF-α. We show here that systemically administered TNF-α was sufficient, alone, to produce acute impairments and is robustly expressed after systemic challenge with LPS (Murray *et al.*, 2012) and after inflammatory trauma such as the tibial fracture used in POCD models (Cibelli *et al.*, 2010; Terrando *et al.*, 2010). The ability of TNF-α to mimic the effects of IL-1 is important in the light of LPS’ propensity to produce equivalent T-maze impairments in wild type and in IL-1R1^-/-^ animals. These IL-1RI^-/-^ animals have developed in the absence of IL-1 signalling (http://jaxmice.jax.org/strain/003245.html) and are known to show normal responses to systemic LPS (Glaccum *et al.*, 1997). Other cytokines, such as TNF-α, demonstrably compensate for the lack of IL-1 signalling to induce sickness behaviour responses in these mice (Bluthe *et al.*, 2000). Reducing systemic cytokines by >90% using DEX (Teeling *et al.*, 2010), here produced robust, though temporally restricted, protection against LPS-induced working memory deficits and TNF-α and IL-1β were both sufficient to mimic LPS effects.

It remains unclear exactly how systemic IL-1 (or TNF-α) alters cognitive function in the current paradigm. However we have already shown that COX-1-mediated prostaglandins contribute to LPS-induced T-maze deficits and ibuprofen was also protective against IL-1-induced deficits (Griffin *et al.*, 2013) so prostaglandins clearly play some causative role. IL-1 may also act directly on vagal afferents or on neurons proximal to the circumventricular organs, which lack a patent blood brain barrier. It is known that peripheral IL-1β can act directly on these neurons and induces expression of the immediate early gene cFOS here and in the amygdaloid complex (Nadjar *et al.*, 2003; Engler *et al.*, 2011), while these neurons do not express TLR4 and are not directly responsive to LPS (Chakravarty & Herkenham, 2005). Similarly IL-1 has robust effects on peripheral energy metabolism (Ota *et al.*, 2009), which may have multiple effects on brain function. These mechanisms require further study.

### Direct effects of IL-1β on hippocampal neurons

Despite equivalent systemic inflammation in NBH and ME7 animals, differential CNS outcomes were observed in ME7 animals and this may occur in several different ways. Firstly, the neurodegenerating brain is ‘primed’ to show exaggerated CNS IL-1β responses to systemic LPS (Cunningham *et al.*, 2005; Godbout *et al.*, 2005; Palin *et al.*, 2008; Pott-Godoy *et al.*, 2008). This exaggerated CNS IL-1 has been assumed to be responsible for the exaggerated sickness behaviour responses to systemic LPS observed in aged and ME7 prion-diseased animals (Combrinck *et al.*, 2002; Godbout *et al.*, 2005) and here we demonstrated that exaggerated hypothermia induced by LPS in ME7 animals is very much mitigated in IL-1R1^-/-^ mice, confirming that IL-1 is a major driver of this heightened sickness response. Secondly, neurons in the diseased brain may be more susceptible to the effects of inflammatory mediators, and concentrations not deleterious to neuronal function in healthy individuals might disrupt function in diseased neurons.

Here we ‘by-passed’ microglial priming by applying equal IL-1β concentrations directly to *ex vivo* hippocampal slices. IL-1β at 0.1 ng/ml (5.9 pM) had no effect on CA1 neurons from NBH animals but was sufficient to depolarize and induce maximal action potential firing in ME7 CA1 neurons. These diseased CA1 neurons were significantly more sensitive (low pM) than prior studies of IL-1-induced depolarization (1 nM: (Coogan & O’Connor, 1997; Ferri & Ferguson, 2003) and the depolarisation observed appeared to be non-synaptic, in that IL-1 had no effect on evoked EPSCs or IPSCs). These effects of IL-1β on CA1 neurons in ME7 animals likely disrupt the precise firing patterns of CA1 pyramidal cells that underlie the rate and temporal codes mediating hippocampal information processing and could contribute to acutely compromised cognitive function in multiple hippocampal-dependent tasks. The *ex vivo* CA1 spiking activity was dependent on IL-1RI^-/-^ and the observation that IL-1β had an irreversible, detrimental effect on CA1 cells from ME7 animals also suggests a role for hippocampal IL-1β in systemic inflammation-induced neuronal death. We have previously shown that increased neuronal death after systemic LPS in ME7 animals was independent of systemic IL-1β (Murray *et al.*, 2011) and in the current study we show that this apoptotic cell death was significantly reduced in IL-1R1^-/-^ mice challenged with LPS. IL-1β applied directly to ME7 neurons was sufficient to produce irreversible depolarization and this effect was IL-1RI^-/-^ dependent. Thus we present evidence for a clear dissociation whereby systemic IL-1β has significant effects on working memory while central IL-1β has significant effects on neuronal viability in the vulnerable brain.

The mechanisms by which IL-1 leads to non-reversible membrane depolarization require further study. IL-1 is widely reported to have pro-convulsant activity and there are data supporting tyrosine kinase phosphorylation of NMDA receptor subunits (Viviani *et al.*, 2003; Balosso *et al.*, 2008), contributing to IL-1-dependent neuronal death. Recently, altered IL-1R1 accessory protein (IL1RAcP) isoform expression in the aged brain has been reported to underpin exaggerated effects of IL-1 in the hippocampus (Prieto *et al.*, 2015) and, while these effects were pertinent to CFC deficits, it is important to investigate whether this mechanism might contribute to decreased viability of neurons in the degenerating brain upon acute elevations of IL-1.

### Conclusion

Systemic IL-1β drives acute working memory dysfunction but centrally acting IL-1β also has robust direct effects on hippocampal and other neuronal populations, leading to hyperexcitation and irreversible loss of membrane potential, and contributing to acute cell death. The data support the idea that the acute and reversible cognitive deficits (including delirium) caused by systemic inflammation may operate via different mechanisms to the concurrent acute brain injury that presumably contributes to the negative long-term cognitive outcomes for patients after recovery from acute cognitive complications such as delirium. Given that delirium contributes to the progression of dementia, and may do so without necessarily exacerbating classical CERAD features like amyloid-β and Tau (Davis *et al.*, 2012; Davis *et al.*, 2017) there are now strong imperatives to understand the extent to which delirium and acute brain injury occur by overlapping or dissociable mechanisms. The relevance of the current model for such investigations has been recently validated with respect to DSM-IV criteria and to alternative models (Davis *et al.*, 2015; Schreuder *et al.*, 2017).

## Acknowledgements

Dr. Colm Cunningham was supported by a Wellcome Trust Senior Fellowship (SRF 090907). DS was supported by a HRB PhD studentship and Edel Hennessy was supported by a Trinity Foundation Studentship. We would like to thank Prof. Kingston Mills for the gift of IL-1R1^-/-^ mice, Stuart Allan for the gift of human IL-1ra and Prof. Roger Anwyl for facilitating electrophysiological studies.

